# Parasympathetic and Sympathetic Activity Are Associated with Individual Differences in Neural Indices of Selective Attention in Adults

**DOI:** 10.1101/173377

**Authors:** Ryan J. Giuliano, Christina M. Karns, Theodore A. Bell, Seth Petersen, Elizabeth A. Skowron, Helen J. Neville, Eric Pakulak

**Affiliations:** Department of Psychology, University of Oregon, Eugene, OR; Prevention Science Institute, University of Oregon, Eugene, OR

**Keywords:** event-related potentials, heart rate variability, pre-ejection period, selective attention, autonomic nervous system, individual differences

## Abstract

Multiple theoretical frameworks posit that interactions between the autonomic nervous system and higher-order neural networks are crucial for cognitive regulation. However, few studies have directly examined the relationship between measures of autonomic physiology and brain activity during cognitive tasks, and fewer studies have examined both the parasympathetic and sympathetic autonomic branches when doing so. Here, 93 adults completed an event-related potential (ERP) auditory selective attention task concurrently with measures of parasympathetic activity (high-frequency heart rate variability; HF-HRV) and sympathetic activity (pre-ejection period; PEP). We replicate previous findings showing effects of selective attention on mean amplitude of the N1 ERP component (Hillyard et al., 1973; Coch et al., 2005), and extend this result to show that the effects of selective attention were associated with baseline values of HF-HRV and PEP. Individuals with higher resting HF-HRV and shorter resting PEP showed larger effects of selective attention on their ERPs. Follow-up regression models demonstrated that HF-HRV and PEP accounted for unique variance in selective attention effects on N1 mean amplitude. These results are consistent with the neurovisceral integration model, which posits that greater parasympathetic activity is a marker of increased cognitive capacity, as well as other theoretical models which emphasize the role of heightened sympathetic activity in more efficient attention-related processing. The present findings highlight the importance of autonomic physiology in the study of individual differences in neurocognitive function, and given the foundational role of selective attention across cognitive domains, suggest that both parasympathetic and sympathetic activity may be key to understanding variability in brain function across a variety of cognitive tasks.

## Introduction

A number of researchers have postulated a critical role for the peripheral nervous system in activity of the central nervous system. Models of neurovisceral integration (Thayer & Lane, 2000; 2009) and polyvagal theory (Porges, 2007) describe similar theoretical frameworks wherein autonomic activity, in particular activity of the parasympathetic nervous system (PNS), is critical for higher-order behavior and cognition due to the high-degree of interconnectedness between neural and peripheral structures regulating behavior and physiological state (Smith, Thayer, Khalsa, & Lane, 2017). However, surprisingly few studies have directly tested associations between autonomic measures and brain activity (Beissner, Meissner, Bar, & Napadow, 2013). Among studies that have examined links between cognitive performance and autonomic physiology, most have focused singularly on measures indexing the PNS (e.g., Thayer & Lane, 2009; Park, Vasey, Van Bavel, & Thayer, 2014; Williams, Thayer, & Koenig, 2016) or sympathetic nervous system (SNS) activity (e.g., Hajcak, McDonald, & Simons, 2003; 2004). Few studies have characterized the joint impact of PNS and SNS function on cognition or underlying neural processes. This is particularly relevant given a wealth of evidence from animal and human studies that autonomic regulation does not operate on a continuum from PNS to SNS dominance, but instead should be conceptualized as interactions between activity in each branch of ‘autonomic space’ (Berntston, Cacioppo, Quigley, 1993b; Berntson, Cacioppo, Quigley, & Fabro, 1994; Levy, 1990). Supporting this framework, we recently found that interactions between PNS and SNS function were associated with individual differences in performance on a difficult working memory task (Giuliano, Gatzke-Kopp, Roos, & Skowron, 2017). Here, we aimed to extend these findings by examining the association between PNS and SNS function and neural mechanisms of selective attention, using a variant of a classic event-related potential (ERP) task designed to control for physiological arousal and minimize psychomotor confounds (Hillyard, Hink, Schwent, & Picton, 1973).

The use of selective attention to examine the interactions between neural mechanisms and autonomic physiology has a number of advantages. First, and primarily, neural mechanisms of selective attention can easily be studied in the absence of manual responses, within contexts where participants are asked to sit calmly and relaxed (e.g., Giuliano, Karns, Neville, & Hillyard, 2014). Motor responses have been shown to be a confound for the interpretation of PNS and SNS measures (Bush, Aikon, Obradovic, Stamperdahl, & Boyce, 2011), yet a majority of the literature demonstrating a relationship between cognitive function and autonomic physiology is based on performance on tasks that require manual, speeded responses (Hansen, Johnsen, & Thayer, 2003; Hansen, Johnsen, Sollers, Stenvik, & Thayer, 2004; Johnsen et al., 2003; Park, Vasey, van Bavel, & Thayer, 2013; Saus et al., 2006; Williams, Thayer, & Koenig, 2016). Second, ERP studies of selective attention have been designed to further control for arousal by measuring effects of attention as the relative difference in the brain response to simultaneously presented to-be-attended and to-be-unattended stimuli (Hillyard, Hink, Schwent, & Picton, 1973). However, it is unclear whether individual differences in physiological measures of arousal influence the pattern of selective attention effects classically observed in this ERP paradigm, nor are we aware of any ERP study of selective attention that has tested PNS and SNS contributions to effects of selective attention. Third, selective attention is a core cognitive skill implicated in a variety of higher-order processes (Cowan et al., 2005; Fukuda, Vogel, Mayr, & Awh, 2010; Garon, Bryson, & Smith, 2008), and the ERP measure of selective attention used here has been shown to predict higher-order cognitive function in preschool-aged children (Isbell, Wray, & Neville, 2015) and in adults (Giuliano, Karns, Neville, & Hillyard, 2014). Therefore, any observed relationship between neural mechanisms of selective attention and autonomic physiology would potentially be relevant for a large body of related cognitive mechanisms. Finally, the ERP selective attention task used here has a high degree of ecological validity and applicability to a wide variety of research samples. Task demands involve sitting still and listening to a narrator within a crowded auditory environment, much like one might be asked to do within an academic or professional context, or even when enjoying leisure time activities. A number of studies have demonstrated that this task is suitable for measuring selective attention from three years of age to adulthood (Karns, Isbell, Giuliano, & Neville, 2015; Sanders. Stevens, Coch, & Neville, 2006), including samples at risk for chronic stress exposure and from lower socioeconomic status backgrounds (Neville et al., 2013; Stevens, Lauinger, & Neville, 2009).

### Parasympathetic nervous system and cognition

Much of the literature on associations between autonomic physiology and cognitive performance is centered on findings that higher resting activity of the PNS often serves as a trait-like marker of regulatory capacity across contexts (Beauchaine & Thayer, 2015; Holzman & Bridgett, 2017). PNS activity is typically quantified as high-frequency heart rate variability (HF-HRV) or respiratory sinus arrhythmia (RSA), both of which are measures of the amount of variability in heart rate that occurs in the respiration bandwidth (Berntson, Cacioppo, & Quigley, 1993a). Activation of the PNS is associated with greater power in the high frequency bandwidth, and serves to quickly decelerate heart rate via projections of the vagal nerve between the brain stem and heart. According to neurovisceral integration theory (Thayer & Lane, 2000; 2009), higher resting levels of HF-HRV or RSA and the associated higher degree of variability in PNS activation reflects a greater capacity for flexible engagement to changes in environmental demands, and is often referred to as an individual’s “vagal tone” (for a commentary, see Berntson, Cacioppo, & Grossman, 2007).

A growing body of evidence suggests that the association between PNS activity and self-regulation extends to domains of cognitive control. For example, on laboratory measures of cognitive ability, adults with higher resting HF-HRV have been shown to have faster and less variable reaction times during target detection tasks (Williams, Thayer, & Koenig, 2016), reaction times that are less affected by the presence of distractors (Park, Vasey, Van Bavel, & Thayer, 2013), faster reaction times during the Eriksen flanker task (Alderman & Olson, 2014), more accurate performance on measures of working memory and sustained attention (Hansen, Johnsen, & Thayer, 2003), and more accurate performance on Stroop tasks in the presence of an additional cognitive load (Capuana, Dywan, Tays, Elmers, Witherspoon, & Segalowitz, 2014). However, the general finding of a positive association between resting HF-HRV activity and performance on cognitive laboratory tasks has a relatively small effect size, with evidence of publication bias towards small significant effects, suggesting this relationship should be interpreted with caution (Zahn, et al., 2016). One likely explanation for why small effect sizes are observed is that resting PNS activity is not associated with cognition broadly, but rather is specifically associated with so-called ‘executive’ processes that emphasize attentional control and inhibition (Kimhy et al., 2013; Spangler, Bell, & Deater-Deckard, 2015). Clarifying the relationship between PNS activity and attentional control processes could have important clinical implications, as individuals with higher resting HF-HRV are more successful at suppressing intrusive thoughts and show related decreases in anxiety and depression symptoms (Gillie, Vasey, & Thayer, 2017).

PNS activity has often been reported to decline or ‘withdraw’ in response to a cognitive challenge (Melis & van Boxtel, 2001; 2007), with an increasing degree of HF-HRV withdrawal reported with increasing task difficulty (Backs & Seljos, 1994; Byrd, Reuther, McNamara, DeLucca, & Berg, 2014; Lenneman & Backs, 2009). For example, there is some evidence that greater HF-HRV withdrawal is associated with faster response times on the Stroop task (Mathewson et al., 2010). However, the directionality of the relationship between PNS reactivity and cognition appears to be context dependent, as increases in HF-HRV activity relative to baseline have been reported in response to challenges that require more regulation of affect (Butler, Wilhelm, & Gross, 2006; Park, Vasey, Van Bavel, & Thayer, 2014; Segerstrom & Nes, 2007).

### Sympathetic nervous system and cognition

In contrast to the importance of resting ‘trait-like’ measures of the PNS for behavior, contributions of the SNS to behavior are typically conceptualized in terms of reactivity. A number of studies have demonstrated that cognitive challenges tend to elicit increases in SNS activation (Allen & Crowell, 1989; Backs & Seljos, 1995; Berntson et al., 1994b; Berntson, Cacioppo, & Fieldstone, 1996). However, two studies by Melis and van Boxtel (2001; 2007) suggest that SNS reactivity is more often seen in association with poor cognitive performance, while PNS reactivity is more relevant to behavior in good performers. In these studies measuring HF-HRV and skin conductance levels during a variety of cognitive tasks, an overall reactivity pattern of HF-HRV withdrawal and increase in skin conductance to the tasks was observed, consistent with previous research. Results from both studies showed that, when comparing good and poor performers, good performers showed a larger effect of PNS withdrawal to the task and had performance that was primarily predicted by PNS withdrawal, while the performance of poor performers was more strongly associated with SNS activation (Melis & van Boxtel, 2001; 2007). These results are supported by findings showing that PNS but not SNS measures are associated with reaction times during a modified Stroop task (Johnsen et al., 2003) and Eriksen flanker task (Alderman & Olson, 2014). It is important to note that many studies of the SNS have relied on measures of galvanic skin response or skin conductance levels, which have been shown to reflect more peripheral fight-or-flight arousal of the electrodermal system (Dawson et al., 2007). Increasingly, studies are employing measures of cardiac pre-ejection period (PEP), a central measure of SNS influences on heart rate derived from the difference in time between the onset of a heart beat (Q point of the QRS complex) and the ejection of blood into the left ventricle, such that greater SNS activation drives shorter ejection intervals (Lozano et al., 2007). PEP has been shown to be more related to reward-sensitive dimensions of the SNS, which include mesolimbic dopaminergic networks also implicated in cardiac regulation (Beauchaine, 2001; Brenner & Beauchaine, 2011). Dissociations between the influence of electrodermal activity and PEP on behavior suggest that electrodermal activity indexes the behavioral inhibition system and avoidance behaviors, while PEP indexes the behavioral activation system and reward-seeking behaviors (Derefinko et al., 2016; Hinnant, Erath, Tu, & El-Sheikh, 2016). Given that reward-sensitivity of mesolimbic dopaminergic processes as indexed by pupillometry has been implicated in individual differences in cognition (Tsukahara, Harrison, & Engle, 2016), it follows that reward-related aspects of the SNS indexed by PEP might be more generally associated with cognition than the avoidance-related aspects of the SNS indexed by electrodermal measures.

### Studies linking neural and autonomic measures

Although a number of meta-analyses have characterized brain structures involved in autonomic activity, there is not a clear consensus on how underlying cortical and subcortical structures give rise to the regulation of PNS and SNS measures. Based on a review of eight neuroimaging studies employing functional MRI and positron emission tomography, Thayer and colleagues (2012) identified the amygdala and ventromedial prefrontal cortex (PFC) as critical regions implicated in PNS regulation (Thayer, Ahs, Fredrikson, Sollers, & Wager, 2012). Within the neurovisceral framework, the authors argue that HF-HRV reflects the degree to which top-down signals emanating from the medial PFC are integrated with brainstem structures regulating cardiac arousal. A separate meta-analysis of 43 neuroimaging studies employing autonomic measures identified significant contributions of both the PNS and SNS to brain activity (Beissner, Meissner, Bar, & Napadow, 2013). Within a subset of eleven studies that measured physiology during cognitive tasks, SNS activity as measured by skin conductance levels was associated with activity in the ventromedial PFC, subgenual anterior cingulate cortex, and amygdala; conversely, PNS activity as measured by HF-HRV was only associated with activity in the anterior insula and amygdala. A separate study of resting brain perfusion similarly found that resting HF-HRV was associated with greater resting blood flow in the anterior insula and amygdala but not in prefrontal regions (Allen, Jennings, Gianaros, Thayer, & Manuck, 2015). Interestingly, a recent theoretical paper emphasizing the role of the PNS for neurovisceral integration does not cite the work by Beissner and colleagues (Smith, Thayer, Khalsa, & Lane, 2017), suggesting that the neurovisceral model could be updated to include evidence documenting the influence of the SNS on activity in midline prefrontal regions during cognitive tasks. Few studies have examined associations between electrophysiological measures of brain activity and measures of autonomic activity. Among the few studies employing ERPs and autonomic measures, Hajcak and colleagues have found that larger ERPs time-locked to response errors are associated with skin conductance measures of increased SNS arousal (Hajcak, McDonald, & Simons, 2003; 2004). Other studies have examined associations between power in different bandwidths of the electroencephalogram (EEG) and autonomic activity. Greater theta power (4-7 Hz) over frontal midline electrodes during an attention-demanding task, an index of attentional control, has been associated with increasing SNS and PNS influences on heart rate (Kubota et al., 2001). At higher bandwidths, asymmetry in alpha power (8-12 Hz) has been associated with SNS measures of skin conductance but not PNS measures of HF-HRV (Gatzke-Kopp, Jetha, & Segalowitz, 2014). Similarly, low-frequency beta power (13-20 Hz) has been reported to decrease with increasing SNS activity, as measured by low-frequency HRV, but was not associated with HF-HRV (Triggiani et al., 2015). As the above review of findings demonstrates, both the SNS and PNS are implicated in measures of brain activity, yet there is still relatively little evidence for associations between autonomic and brain activity with regard to individual differences in performance.

### The current study

Here we aimed to characterize PNS and SNS function during a selective attention task and clarify how PNS and SNS activity relates to concurrent neural measures of selective attention. We sought to establish these relationships within a family context using a sample of parents from a larger study involving children and their caregivers. Our community sample of parents, primarily mothers, is advantageous in that similar samples are frequently targeted for educational and health interventions (Shonkoff & Fisher, 2013), and is consistent with recommendations for psychophysiological studies to be performed beyond a university population (Gatzke-Kopp, 2016).

Adult participants volunteered for a laboratory visit where HF-HRV and PEP were measured as indices of PNS and SNS function, respectively, during a 5-minute neutral film clip and subsequently during four separate stories of an auditory attention task designed for the recording of ERPs. Analyses focused on modulations of selective attention on the ERP elicited by identical sound probes presented at to-be-attended and to-be-ignored spatial locations, and associations between modulations of selective attention on ERPs with measures of HF-HRV and PEP at baseline and during the task. Based on previous research, we hypothesized that higher baseline HF-HRV would be associated with larger ERP effects of selective attention. Additionally, we hypothesized that greater SNS arousal, as indexed by shorter PEP, would be associated with larger ERP effects of selective attention. To the extent that the task elicited reactivity in HF-HRV and PEP values relative to baseline, we hypothesized that greater reactivity would be associated with larger effects of selective attention, consistent with the view that greater flexibility in physiological activity is associated with better cognitive function (Thayer & Lane, 2009).

## Method

### Participants

Adults were recruited along with their children as part of a broader study testing a two-generation training program integrated into Head Start services. Adults were eligible to participate in laboratory visits if they were English-language dominant, had no history of neurological or hearing impairment, and normal or corrected-to-normal vision. Electroencephalogram (EEG) measures were successfully obtained from 129 adults. Seven adults declined placement of electrodes for measurement of electrocardiogram (ECG), eighteen adults did not have appropriately event-marked ECG data, and eleven adults had unusable impedance cardiogram (IC) data at baseline (n = 4), during the task (n = 4), or both (n = 3). This left 93 adults with complete data (87 females; age, *M* = 32.56 years, *SD* = 7.51, range = 32 to 67 years). The majority of the adults recruited were female, so all analyses were duplicated without the six male participants and are reported in the supplementary results (see Supplemental Tables 1 and 2).

Adult participants enrolled in this study were all parents of a child currently enrolled in Head Start, and thus were at increased risk for living at or below the poverty line. To quantify the degree of exposure to risk factors related to lower socioeconomic status, we calculated a cumulative index of socioeconomic risk based on three discrete, additive markers: household income, maternal education, and maternal marital status (Evans & Kim, 2013). For each of the three indicators, risk was coded dichotomously as present (1) or not present (0), with each risk factor aimed to characterize the riskiest third of the sample for that dimension. Household income risk was quantified as households in the bottom third of annual incomes within the sample, here annual incomes less than $15,000 (N=37). Maternal education risk was quantified as households in which the mother’s highest degree of education was a high school diploma or less (N=30). Marital status risk was quantified as single parent households, in this case all single mothers (N=30). Based on this additive metric of socioeconomic risk, zero risks were present for 30 families, one risk was present for 36 participants, and two or three risks were present for 27 participants. As noted in the results below, analyses examining associations between cumulative socioeconomic risks and the individual risk factors separately showed no associations with ERPs elicited during the selective attention task or any of the measures of autonomic physiology.

### Measures

#### Auditory selective attention task

We used the same auditory attention ERP paradigm as our previous studies with child and adult participants (Karns, Isbell, Giuliano, & Neville, 2015; Neville et al., 2013; Stevens et al., 2009; Sanders et al., 2006; Coch et al., 2005). Participants were cued to selectively attend to one of two simultaneously presented children’s stories differing in location (left/right loudspeaker), narrator’s voice (a male or female reading the entire story aloud), and content. Illustrations from the story being read from the attended loudspeaker were presented on a monitor. A green arrow pointing to the left or right was displayed on the monitor throughout each block to indicate the side of the to-be-attended story.

ERPs were recorded to 100 ms duration probe stimuli embedded in both to-be-attended and to-be-unattended stories. Probe stimuli were either linguistic (a voiced syllable) or nonlinguistic (a broad-spectrum buzz). The linguistic probe was the syllable /ba/, spoken by a female (a different voice from all the story narrators). The nonlinguistic probe was created by scrambling 4-6ms segments of the /ba/ stimulus, which preserved many of the acoustic properties of the linguistic probe. Across the stories, approximately 200 linguistic and 200 nonlinguistic probes were presented in each channel. The probes were presented randomly at an inter-stimulus interval (ISI) of either 200, 500, or 1000ms in one of the two loudspeakers at a time, with a range of plus or minus 25 ms of jitter for each ISI. The stories were presented at an average of 60 dB SPL, and the intensity of the probes was 70 dB. A researcher monitored adults throughout the task to ensure that they remained still and equidistant between the two loudspeakers, and to administer comprehension questions via intercom following each pair of simultaneously presented stories.

Within each testing session, a total of eight different stories were presented. Since two stories were presented simultaneously, there were a total of four listening blocks. Each listening block ranged in length from four to six minutes long resulting in approximately 20 minutes of on-task time, with approximately one to two minutes of off-task time between blocks. The stories were selected from the following children’s book series: *Blue Kangaroo* (Clark, 1998, 2000, 2002, 2003), *Harry the Dog* (Zion & Graham, 1956, 1960, 1965, 1976), Max & Ruby (Wells, 1997, 2000, 2002, 2004), and *Munsch for Kids* (Munsch & Martchenko, 1989, 2001, 2002; Munsch & Suomalainen, 1995). In each of the four blocks, two stories were presented simultaneously with attention directed to the story played from either the left or right loudspeaker and read aloud by a different narrator from four narrators in total. For example, in one block, a participant could be directed to listen to the left loudspeaker, which might play a story read by a female narrator, while ignoring the right loudspeaker, which would be playing a story read by a male narrator; the illustrations on the monitor would correspond to the attended story. Each participant attended twice to a story on the right side and twice to a story on the left side, with start side counterbalanced across participants. For each participant, an individual narrator would be heard once as the narrator for an attended story and once as the narrator for an unattended story. After each story, an experimenter asked the participant three open-ended comprehension questions about the attended story to reinforce the goal of paying attention. The comprehension questions were written to emphasize multiple pieces of information that occurred throughout the entire course of the story. Verbal responses were coded by the experimenter, who marked any keywords of the true answer spoken by the participant when answering the question.

#### Electroencephalogram (EEG)

During the dichotic listening task, EEG was recorded at a sampling rate of 512 Hz from 64 Ag/Ag-Cl-tipped scalp electrodes (BioSemi Active2, Amsterdam, Netherlands) arranged according to the international 10–20 system. Electrode offsets were maintained at ±30 μV or less throughout each recording session. Additional electrodes were placed on the outer canthus of each eye, below the right eye, and on the left and right mastoids. The EEG was recorded relative to the common mode sense active electrode and then re-referenced offline to the algebraic mean of the left and right mastoids. Horizontal eye movements were plotted as the difference between the left and right outer canthus channels. Vertical eye movement, including eye blinks, were plotted as the difference between the lower right eye electrode and Fp1 (left anterior-most electrode). ERP analyses were carried out using EEGLAB (Delorme & Makeig, 2004) and ERPLAB (Lopez-Calderon & Luck, 2014). Raw EEG data were imported into EEGLAB and high-pass filtered at 0.1 Hz. Then, epochs time-locked to sound probes embedded in the dichotic listening task were extracted from -100 to 500 msec relative to probe onset. Epochs containing large voltage deviations or muscle/movement artifacts were identified by visual inspection and excluded from further analysis. Remaining data were then submitted to artifact rejection procedures within ERPLAB. Artifacts were identified based on moving window peak-to-peak changes in eye channels across a 200 ms window, moving in 50 ms increments. Individual artifact rejection parameters were adjusted for each participant to identify the smallest amplitude changes associated with eye movements or blinks. Manual artifact rejection was employed after the automatic ERPLAB procedures, to ensure accuracy of artifact marking. Following artifact rejection, epochs were low-pass filtered at 40 Hz.

In order to facilitate analyses of the distribution of ERP effects and to reduce the number of multiple comparisons across electrode sites, we averaged the 64 electrodes into nine electrode clusters representing the scalp with distributional factors in a 3x3 design of anteriority (anterior, central, posterior) x laterality (left, medial, and right). The electrode clusters consisted of Left Anterior (AF3, AF7, F3, F5, F7), Medial Anterior (AFz, Fz, F1, F2), Right Anterior (AF4, AF8, F4, F6, F8), Left Central (FC3, FC5, C3, C5, FT7, T7), Medial Central (FCz, Cz, FC1, FC2, C1, C2), Right Central (FC4, FC6, C4, C6, FT8, T8), Left Posterior (CP3, CP5, P3, P5, P7, PO3, PO7, TP7), Medial Posterior (CPz, Pz, POz, CP1, CP2, P1, P2), and Right Posterior (CP4, CP6, P4, P6, P8, PO4, PO8, TP8). Mean amplitudes were extracted from time-windows of interest and subjected to an attention (2) x anteriority (3) x laterality (3) repeated-measures ANOVA to examine effects of selective attention. We note that in order to reduce the number of interactions being tested for, this statistical model collapses across probe type by averaging together the mean amplitude of ERP responses to linguistic and nonlinguistic sound probes. At time windows where there were significant effects of attention on ERP amplitudes, we examined associations between ERPs, HF-HRV, and PEP. In order to reduce the number of multiple comparisons, we averaged across electrode clusters demonstrating significant effects of attention, then examined partial correlations between ERP amplitudes and baseline HF-HRV, HF-HRV reactivity, baseline PEP, and PEP reactivity. Significant associations were followed with an examination of partial correlations with attended and unattended ERPs separately. Then, linear regression models were run to examine significant relationships between ERP effects of attention and physiological measures when including all four physiological measures in the model, with stepwise entering of age, then baseline HF-HRV and baseline PEP, followed by HF-HRV reactivity and PEP reactivity. After these primary ERP analyses, we then explored correlations between ERPs and HF-HRV and PEP across the scalp, including correlations with socioeconomic risk levels^1^.

#### Cardiovascular physiology

A montage of 11 electrodes was used for the measurement of high-frequency heart rate variability (HF-HRV) and pre-ejection period (PEP). Electrocardiogram (ECG) was obtained from three disposable pre-gelled electrodes placed in a modified Lead II configuration on the distal right clavicle, lower left rib, and lower right abdomen. Impedance cardiogram (ICG; Z0) was recorded from eight electrodes placed in a tetrapolar configuration on the left and right lateral neck and torso, from a vertical maximum of the jawline down to the diaphragm. Data were acquired wirelessly via Biopac Nomadix BN-RSPEC and BN-NICO transmitters (Biopac Systems Inc, Goleta, CA) sending ECG and impedance signals respectively to a Biopac MP-150 acquisition unit placed in the room with the participant. A respiration signal was derived from the raw impedance cardiogram for the inspection of respiration values. High-frequency HF-HRV values were derived from natural log-transformed values of the spectral power in the high frequency range commonly used for adults (.12-.40 Hz). PEP was calculated from the first-order derivative of the cardiovascular impedance signal (dZ/dt), as the length of time from the Q-point of the ECG waveform to the B-point of the dZ/dt waveform (Berntson, Lozano, Chen, & Cacioppo, 2004).

Autonomic data were processed separately for a 5-minute baseline period and for the four blocks of the dichotic listening task. Data processing was performed using Mindware HF-HRV and IMP softwares (Gahanna, OH). First, ECG signals were inspected by trained research assistants to ensure the correct identification of individual R peaks. Edited ECG files were then used for the processing of PEP values, whereby visual inspection was used to verify that both the Q- and B-points were present and correctly placed in 30-second averages of ECG and dz/dt waveforms. HF-HRV, PEP, heart rate, and respiration rate values were exported in 30-second epochs, then averaged across epochs to yield a single baseline value and a separate value for each of the four blocks of the dichotic listening task. Physiological reactivity values during the task were calculated for all measures as difference scores from baseline values (task minus baseline). For HF-HRV, positive reactivity scores index greater PNS activation and negative scores index PNS withdrawal, relative to baseline levels. Because longer PEP intervals reflect less SNS activation, positive PEP change scores reflect SNS withdrawal during the task, and negative PEP change scores reflect SNS activation during the task, relative to baseline.

### Procedure

Upon arrival to the laboratory, informed consent was obtained and electrode application was initiated. First, electrodes for the recording of ECG and IC were placed onto the participant’s torso, followed by placement of an electrode cap and electrodes for EEG recording. During this time, electrodes for monitoring of ECG, IC, and EEG were also being placed onto the participant’s child (Giuliano et al., *under review*). Then, the participant and child were ushered into an electrically-shielded, sound-attenuating booth for the baseline physiology recording. Children were seated in a comfortable chair positioned 145 cm away from a computer monitor, with two speakers placed 90° to the left and right of the chair, while parents were seated in a chair to the right of the child. Then, an initial five-minute “baseline” measure of ECG and IC were taken while a video depicting calming ocean scenes and featuring low volume instrumental music was presented on the monitor. After the ocean video completed, a research assistant entered and ushered the parent out and into another booth down the hallway. A research assistant followed the parent into the new booth for initiation of the EEG recording, where parents were seated in a chair positioned 145 cm away from a computer monitor, with two speakers placed 90° to the left and right of the chair. Participants first heard instructions, followed by four stories with three open-ended comprehension questions asked in between each story to ensure participant attention to the task. After completion of the task, participants moved on to complete additional testing procedures (not reported here).

## Results

### Characterizing physiological reactivity to the selective attention task

Descriptive statistics for all physiological measures are shown in Table 1 below. Twotailed paired-sample *t*-tests of baseline and task values for HF-HRV, PEP, heart rate, and respiration rate showed significant task reactivity for all measures. HF-HRV power declined from baseline (*M* = 6.09, *SD* = 1.13) to task (*M* = 5.98, *SD* = 1.06), *t*(92) = 2.01, *p* = .047. PEP values shortened from baseline (*M* = 113.93, *SD* = 10.95) to task (*M* = 112.88, *SD* = 10.65), *t*(92) = 2.02, *p* = .047. These changes in HF-HRV and PEP were associated with concurrent slowing of heart rate from baseline (*M* = 73.15, *SD* = 10.27) to task (*M* = 71.87, *SD* = 9.66), *t*(92) = 3.23, *p* = .002, and acceleration of respiration rate from baseline (*M* = 15.84, *SD* = 1.82) to task (*M* = 17.35, *SD* = 2.36), *t*(92) = -7.56, *p* < .001.

**Table 1.** Descriptive Statistics.

|  | *M* | *SD* | Range |
| --- | --- | --- | --- |
| Participant age | 32.56 | 7.51 | 22–67 |
| SES risk index | 1.02 | 0.88 | 0–3 |
| Baseline HF-HRV | 6.09 | 1.13 | 3.28–8.52 |
| Task HF-HRV | 5.98 | 1.06 | 2.97–8.36 |
| HF-HRV reactivity | -0.11 | 0.50 | -2.12–1.15 |
| Baseline PEP | 113.93 | 10.95 | 87.80–143.60 |
| Task PEP | 112.88 | 10.65 | 87.82–139.61 |
| PEP reactivity | -1.05 | 5.02 | -10.81–16.15 |
| Baseline heart rate | 73.15 | 10.27 | 50.21–103.97 |
| Task heart rate | 71.87 | 9.66 | 52.23–97.18 |
| Heart rate reactivity | -1.27 | 3.81 | -9.89–8.49 |
| Baseline respiration rate | 15.84 | 1.82 | 11.70–21.66 |
| Task respiration rate | 17.35 | 2.36 | 12.18–24.55 |
| Respiration rate reactivity | 1.51 | 1.92 | -4.40–8.33 |

### Characterizing effects of selective attention on ERPs

Visual inspection of ERP waveforms elicited by sound probes embedded in attended and unattended stories can be seen in Figure 2 below. For both probe conditions, a frontocentral maximum positive deflection can be seen peaking after 100 ms (“P1”), followed by a phasic negative deflection peaking before 200 ms (“N1”). Statistical analyses were performed separately on mean amplitudes extracted from ERPs in 50 ms windows centered on the P1 and N1 peaks, 100-150 ms and 175-225 ms respectively. For P1 mean amplitudes, results revealed an interaction of attention × laterality, *F*(2, 184) = 3.41, *p* = .037, such that significant effects of attention were observed at right-lateralized electrode clusters (*p* = .020) but not at left-lateralized or midline clusters (*ps* > .35). Follow-up comparisons showed that significant effects of attention on P1 amplitudes at the group level were observed at the right medial (*p* = .009) and right posterior (*p* = .037) electrode clusters. For N1 mean amplitudes, results revealed a main effect of attention, *F*(1, 92) = 7.25, *p* = .008, as well as an interaction of attention x laterality, *F*(2, 184) = 11.36, *p* < .001, such that significant effects of attention were observed at left-lateralized (*p* = .007) and midline electrode clusters (*p* < .001) but not at the right-lateralized clusters (*p* = .553). Follow-up comparisons demonstrated significant attention effects on N1 amplitudes broadly across the scalp [left anterior, *p* = .027; central anterior, *p* = .012; central midline, *p* = .001; left posterior, *p* = .003; midline posterior, *p* < .001]. Therefore, subsequent analyses quantified P1 amplitudes as a composite of the right medial and right posterior electrode clusters, while N1 amplitudes were quantified as a composite of the left anterior, midline anterior, midline central, left posterior, and midline posterior electrode clusters.

**Figure 1.**
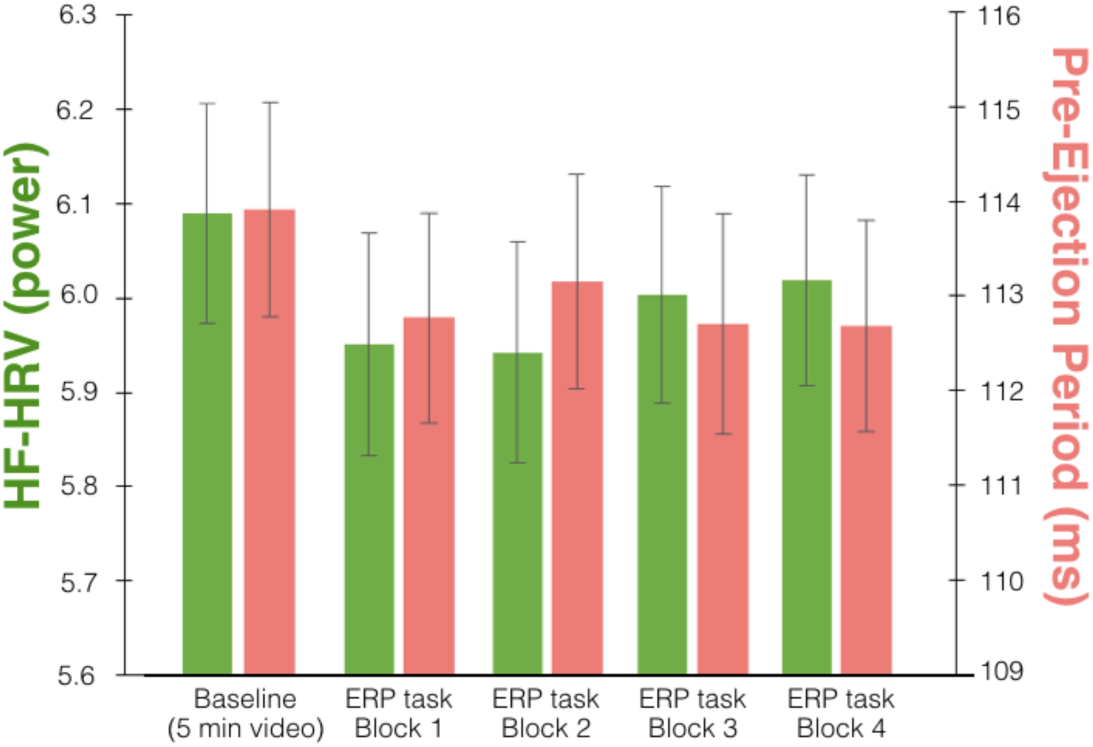
HF-HRV and PEP values during baseline and four stories of the selective attention task. Standard error bars are shown.

**Figure 2.**
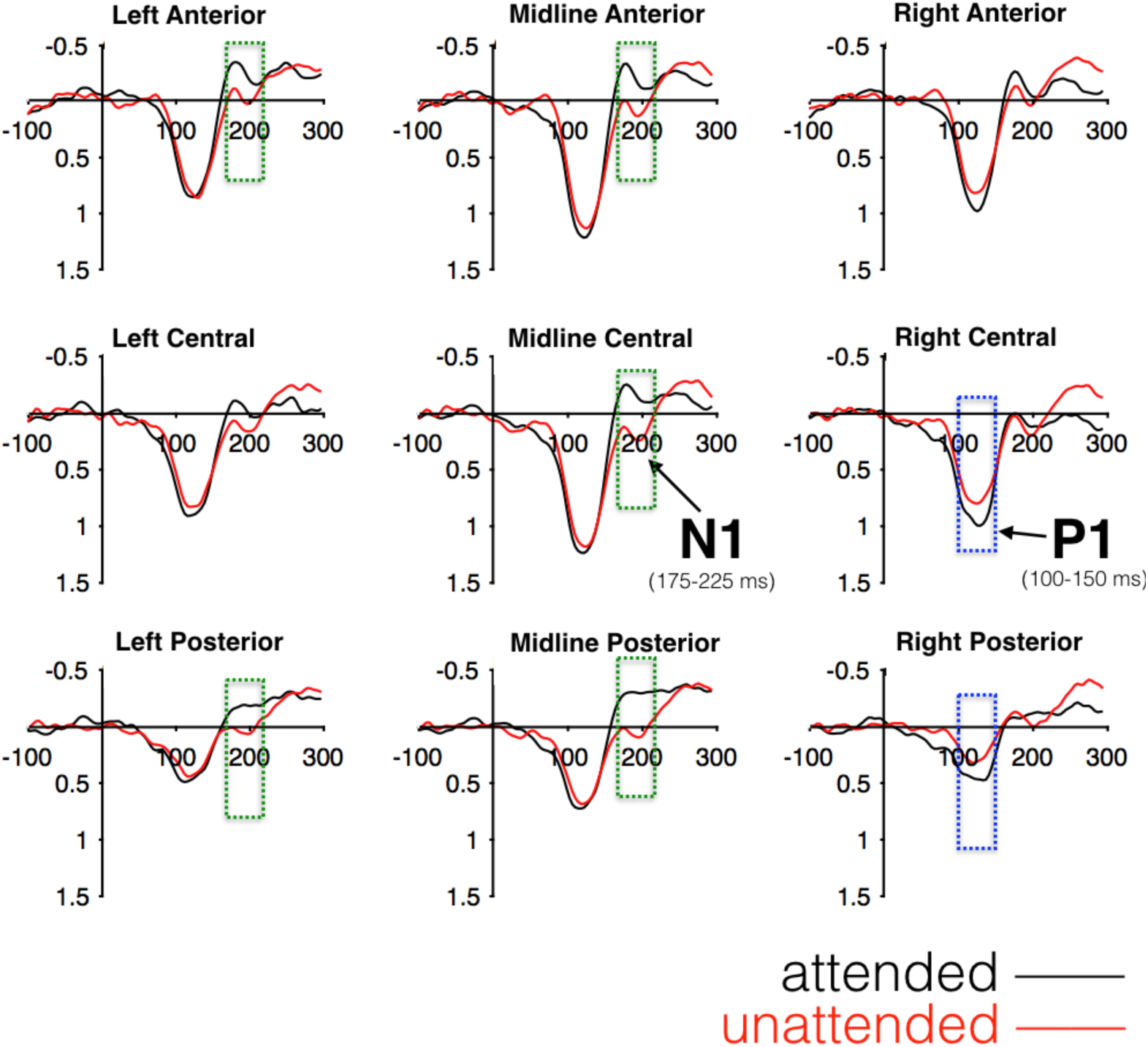
ERPs to attended and unattended probes for 93 adult participants. Electrode clusters highlighted in blue were averaged into a composite for P1 analyses, and clusters in green were averaged into a separate composite for N1 analyses.

### Associations between ANS and ERP effects of selective attention

Partial correlations among all variables of interest while controlling for participant age are shown in Table 2. Below, these results and follow-up analyses are presented separately for each ERP component. Notably, there were no significant associations between socioeconomic risk factors and any of the autonomic or ERP measures. Therefore, socioeconomic risk was not included in the following analyses.

**Table 2.** Partial Correlations Among All Variables of Interest While Controlling for Age.

|  | 1 | 2 | 3 | 4 | 5 | 6 | 7 | 8 | 9 | 10 | 11 | 12 |
| --- | --- | --- | --- | --- | --- | --- | --- | --- | --- | --- | --- | --- |
| 1. Socioeconomic risk | - |  |  |  |  |  |  |  |  |  |  |  |
| 2. HF-HRV baseline | .06 | - |  |  |  |  |  |  |  |  |  |  |
| 3. HF-HRV task | -.03 | .88*** | - |  |  |  |  |  |  |  |  |  |
| 4. HF-HRV reactivity | -.17 | -.34** | .15 | - |  |  |  |  |  |  |  |  |
| 5. PEP baseline | -.03 | .06 | .11 | .09 | - |  |  |  |  |  |  |  |
| 6. PEP task | -.09 | .04 | .07 | .07 | .90** | - |  |  |  |  |  |  |
| 7. PEP reactivity | -.13 | -.05 | -.08 | -.06 | .30** | .16 | - |  |  |  |  |  |
| 8. Attended P1 amp. | .06 | -.05 | -.10 | -.10 | .17 | .07 | -.22* | - |  |  |  |  |
| 9. Unattended P1 amp. | .17 | -.15 | -.15 | .03 | .08 | .04 | -.09 | .32** | - |  |  |  |
| 10. Att.-Unatt. P1 amp. | -.09 | .08 | .02 | -.12 | .08 | .03 | -.12 | .62*** | -.54*** | - |  |  |
| 11. Attended N1 amp. | .04 | -.19t | -.16 | .08 | .25* | .29** | .07 | .03 | <.01 | .03 | - |  |
| 12. Unatt. N1 amp. | .10 | .03 | .02 | -.02 | -.01 | -.02 | -.02 | -.16 | .20t | -.30** | .46*** | - |
| 13. Att.- Unatt. N1 amp. | -.06 | -.22* | -.18 | .10 | .26* | .31** | .08 | .17 | -.18 | .30** | .57*** | -.46*** |

Note: \*\*\*, $p < .001$ ; \*\*, $p < .01$ ; \*, $p < .05$ ; t, $p < .07$

### P1 component (100-150 ms)

Partial correlations controlling for age revealed no significant associations between effects of attention on P1 amplitudes and measures of HF-HRV [baseline HF-HRV: *r*(90) = .08, *p* = .459; task HF-HRV: *r*(90) = .02, *p* = .822; HF-HRV reactivity: *r*(90) = −.12, *p* = .271] or measures of PEP [baseline PEP: *r*(90) = .08, *p* = .425; task PEP: *r*(90) = .03, *p* = .767; PEP reactivity: *r*(90) = −.12, *p* = .255].

### N1 component (175-225 ms)

Partial correlations controlling for age revealed significant associations between effects of selective attention on N1 amplitudes and both baseline HF-HRV, *r*(90) = −.22, *p* = .037, and baseline PEP, *r*(90) = .26, *p* = .011. A similar effect was observed between N1 amplitudes and task values of PEP, *r*(90) = .31, *p* = .003, although there was only a trend of N1 amplitudes correlating with task HF-HRV, *r*(90) = −.18, *p* = .086. As seen in Figure 2.3, higher baseline HF-HRV and shorter baseline PEP were associated with a larger effect of selective attention on N1 amplitudes. Notably, these effects remained when controlling for respiration rate [baseline HF-HRV, *r*(90) = −.21, *p* = .046; baseline PEP, *r*(90) = .26, *p* = .013]^2^. Follow-up analyses of attended and unattended ERPs separately indicated that the relationship between baseline physiology and the N1 attention effect was driven by associations specific to attended ERPs. Shorter baseline PEP was associated with larger negative amplitudes to attended ERPs, *r*(90) = .25, *p* = .015, and a similar trend was observed between higher baseline HF-HRV and larger negative amplitudes to attended ERPs, *r*(90) = −.19, *p* = .066. There was no evidence of associations between baseline physiology and unattended ERPs (*ps* > .79).

**Figure 3.**
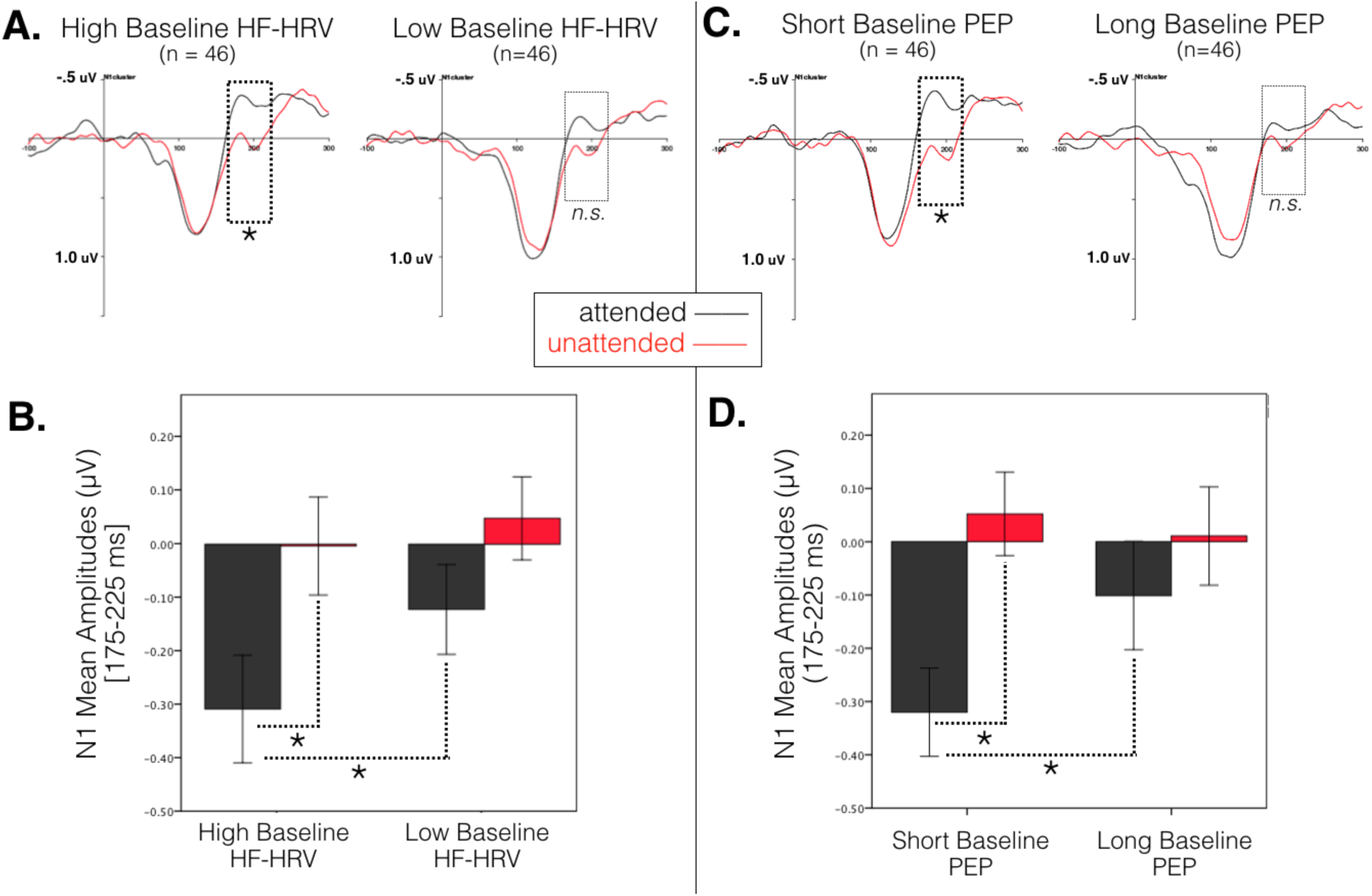
Composite ERPs aggregated across frontocentral electrode sites used in N1 analyses presented by median split on baseline HF-HRV and baseline PEP values. Note that analyses were conducted with continuous measures. A) ERPs illustrated by median split on baseline HF-HRV. B) Mean amplitudes of the N1 component for attended and unattended probes by median split on baseline HF-HRV. C) ERPs illustrated by mean split on baseline PEP. D) Mean amplitudes of the N1 component for attended and unattended probes by median split on baseline PEP.

Linear regression models were then examined to clarify the joint contributions of baseline HF-HRV and baseline PEP with effects of selective attention on N1 amplitudes, while controlling for reactivity values and age. As shown in Table 3, the effect of attention on N1 amplitudes was predicted by baseline HF-HRV and baseline PEP [*R*^2^ = .14, *F*(3, 89) = 4.80, *p* = .004], with additional variance accounted for by baseline HF-HRV [*β* = -.25, *p* = .021] and baseline PEP [*β* = .27, *p* = .007]. Adding reactivity values of HF-HRV and PEP did not contribute additional explained variance to the model [*R*^2^ change = .02, *F* change(2, 87) = 1.26, *p* = .289]. To test for interactions between baseline HF-HRV and PEP, an additional model was performed including an interaction term of baseline HF-HRV x baseline PEP, but adding the interaction term did not contribute additional explained variance [*R*^2^ change = .02, *F* change(1, 86) = 1.80, *p* = .183]. In summary, the N1 attention effect showed a negative correlation with baseline HF-HRV and a positive correlation with baseline PEP: larger attention effects on the N1 component were associated with both greater baseline HF-HRV and shorter baseline PEP.

**Table 3.** Regression Models Predicting the Effect of Selective Attention on N1 Amplitudes from HF-HRV and PEP values.

| Model 1 |  |  | Model 2 |  |  | Model 3 |  |  |
| --- | --- | --- | --- | --- | --- | --- | --- | --- |
| | $\beta$ | $p$ | | $\beta$ | $p$ | | $\beta$ | $p$ |
| Age | .03 | .766 | Age | <.01 | .977 | Age | -.01 | .896 |
| **Baseline PEP** | **.27** | **.007\*\*** | **Baseline PEP** | **.32** | **.002\*\*** | **Baseline PEP** | **.31** | **.004\*\*** |
| **Baseline HF-HRV** | **-.25** | **.021\*** | **Baseline HF-HRV** | **-.24** | **.033\*** | **Baseline HF-HRV** | **-.26** | **.026\*** |
|  |  |  | PEP reactivity | .17 | .117 | PEP reactivity | .16 | .126 |
|  |  |  | HF-HRV reactivity | <.01 | .996 | HF-HRV reactivity | .02 | .856 |
|  |  |  |  |  |  | Baseline PEP x HF-HRV | -.13 | .183 |
| Model fit, $F(3, 89) = 4.80$, $p = .004**$<br>$R^2 = .139$ | | | Model fit, $F(5, 87) = 3.40$, $p = .007**$<br>$F$ change (2, 87) = 1.26, $p = .289$<br>$R^2 = .163$ | | | Model fit, $F(6, 86) = 3.16$, $p = .008**$<br>$F$ change (1, 86) = 1.80, $p = .183$<br>$R^2 = .181$ | | |

As an additional control, we examined whether associations between ERPs and autonomic physiology were confounded by the number of artifact-free trials available for ERP averaging. Zero-order correlations demonstrated a significant negative correlation between the number of artifact-free trials and baseline HF-HRV, *r*(91) = -.33, *p* = .001, such that higher baseline HF-HRV was associated with fewer artifact-free trials. No relationships were observed between artifact-free trials and HF-HRV reactivity (*p* = .387) or any measure of PEP (all *p*s > .31). Partial correlations between the N1 attention effect and baseline HF-HRV and baseline PEP were examined, while controlling for number of artifact-free trials. Significant relationships were still observed between the N1 attention effect and baseline HF-HRV, *r*(90) = -.21, *p* = .042, and between the N1 attention effect and baseline PEP, *r*(90) = .29, *p* = .006.

## Discussion

This study sought to characterize the extent to which ERP mechanisms of selective attention are associated with markers of parasympathetic and sympathetic function, specifically HF-HRV and PEP, in adults from a community sample with children in Head Start. We observed a widely-distributed effect of selective attention on ERP amplitudes at the N1 component that was associated with baseline levels of HF-HRV and PEP. As predicted, individuals with higher baseline HF-HRV and shorter baseline PEP showed larger effects of selective attention. No relationship between current SES and autonomic physiology or ERPs was observed. These results demonstrate the importance of both autonomic branches for neural activity and cognitive performance, such that physiological components of flexible engagement and reward-sensitivity may be independent dimensions contributing to optimal neurocognitive function.

Consistent with a number studies within the neurovisceral integration framework demonstrating a positive association between resting PNS activity and cognitive measures of attention (for a review, see Park & Thayer, 2014), greater baseline HF-HRV was associated with larger effects of selective attention on ERPs. Greater variability in high-frequency bandwidth of heart rate at rest has been proposed to reflect a greater capacity for self-regulation and cognitive performance, and underlying activity in the prefrontal cortex related to goal-directed behavior (Smith, Thayer, Khalsa, & Lane, 2017). Indeed, lesions to the prefrontal cortex have been associated with marked attenuations of the N1 attention effect in the selective attention used here (Knight, Hillyard, Woods, & Neville, 1981), suggesting that individuals with lower resting HF-HRV in the current study might also be individuals who exhibit less goal-directed activity (e.g. top-down control signals) in networks involving the prefrontal cortex.

Larger selective attention effects on ERPs were also associated with shorter resting PEP intervals, consistent with previous studies finding positive associations between increased SNS activity and brain function (Beissner et al., 2013; Hajcak et al., 2003; 2004). In a review of autonomic influences on attention performance, Hugdahl noted that an increased galvanic skin response often accompanies sustained orienting and habituation to repeated stimuli, particularly with regard to demands for attentional control (Hugdahl, 1996). Interestingly, a similar relationship between increased galvanic skin response and higher-order cognitive processes has been proposed by several researchers (Edelberg, 1993; Tranel & Damasio, 1994; Ohman, 1992). In particular, given that lesions to the prefrontal cortex have been associated with an elimination of the galvanic skin response to attention-demanding stimuli (Tranel & Damasio, 1994), the SNS seems similarly important for selective attention as the PNS. Yet, many reports of relationships between autonomic activity and attention only report measures of HF-HRV or heart rate (e.g., Park et al., 2013; 2014; Williams et al., 2016), suggesting that interpretations of the relationship between autonomic activity and attention may be biased towards PNS contributions to attention. For example, interpretations from studies employing PNS-mediated measures such as HF-HRV have emphasized the role of top-down regulation in attentional control (Park & Thayer, 2014), while interpretations based on studies employing SNS measures such as galvanic skin response have emphasized the role of bottom-up orienting responses (e.g., Hugdahl, 1996).

A consideration of the simultaneous contributions of resting HF-HRV and PEP to effects of selective attention observed here lends support for both views of autonomic contributions to attention, but leads to a more nuanced interpretation than would have likely been reached from either measure either autonomic system in isolation. The finding that higher resting HF-HRV is associated with better selective attention supports literature suggesting these same adults with higher HF-HRV have a greater capacity for regulation of behavior and attention. One interpretation of why this relationship exists might be that adults with higher resting PNS activation are calmer when they start the attention task, and that calm state may facilitate better selective attention to the stories of interest within the crowded environment. However, this interpretation seems unlikely when considering PEP, as larger attention effects were associated with shorter resting PEP, or greater sympathetic activation. A similarly short-sighted conclusion might be drawn from considering PEP in isolation, such that a more aroused and SNS-activated state might be seen as better for selective attention performance. Considering both autonomic systems in tandem leads to a more refined interpretation: given evidence that shorter PEP reflects greater reward-related SNS activity as opposed to threat-related SNS activity (e.g., Brenner & Beauchaine, 2011), the co-occurence of high resting HF-HRV and shorter resting PEP may reflect a general disposition of high selfregulation in the context of high approach-related behaviors. It is still possible that greater threat-related SNS arousal is indeed related to better selective attention performance, but this result must still be considered within the context of heightened PNS activity showing a similar relationship with attention. Given previous evidence that optimal behavioral performance is typically observed with reciprocally-activating systems (Melis & van Boxtel, 2001; 2007), the interpretation consistent with reciprocal activity suggests the shorter PEP observed here in the context of higher HF-HRV is due to shorter PEP indexing greater reward-related approach to the laboratory visit. This interpretation is also consistent with the experimental context of the selective attention task used here, where participants are asked to sit calmly without exposure to additional stressors or threats.

One limitation to the interpretation of the present findings is that the baseline physiological measurement was taken while adults were seated with their child in the ERP booth. While joint parent-child baseline measurements are often used in studies involving participants who are parents (e.g. Giuliano, Skowron, & Berkman, 2015), it is possible that adults in this study were actively engaged in self-regulation and parenting during the baseline measurement in order to facilitate their children sitting still and quiet through the 5-minute baseline video. If so, the baseline measure of HF-HRV may capture some degree of resting HF-HRV levels in addition to augmented HF-HRV to the extent that individuals were engaged in regulated parenting (Skowron, Cipriano-Essel, Benjamin, Pincus, & Van Ryzin, 2013). Although less is known about PEP reactivity in the context of parenting, baseline PEP values may be similarly biased.

Another limitation to the present study concerns the generalizability of the findings based on a community sample of adults living at risk for a variety of poverty-related stressors. However, there is already a large body of research on higher SES populations, and researchers have called for studies with more diverse samples in psychophysical studies (Gatzke-Kopp, 2016). This sample is of particular relevance given efforts of translational science to harness the neuroplasticity of selective attention mechanisms in evidence-based interventions targeting these systems (e.g., Neville et al., 2013). Thus, it is critical to characterize the relationship between attention and measures of autonomic physiology in individuals who would most likely be recruited for such family training programs. Even if the associations between autonomic physiology and selective attention reported here are unique to a population living in lower SES contexts, this information is still relevant as evidence for intervention science, and suggests that a consideration of influences on autonomic physiology when designing intervention strategies might facilitate efforts to train selective attention. Future research with adults from a wide range of socioeconomic backgrounds during childhood, in addition to better characterizing a variety of stressful life events, will shed further light on how early experience may impact the relationship between autonomic and neural mechanisms during selective attention.

The lack of correlations between measures of HF-HRV and PEP observed here replicates a wide body of work demonstrating that the responses of the two autonomic branches represent separate dimensions of cardiac regulation (Berntson, Cacioppo, & Quigley, 1993b; Berntson et al., 1994; Berntson, Cacioppo, & Fieldstone, 1996). Consistent with our previous work, the two autonomic branches as indexed by HF-HRV and PEP appear to make unique contributions to cognitive function (Giuliano, Gatzke-Kopp, Roos, & Skowron, 2017). What remains to be seen is to what extent cognitive processing is related to reward-related versus threat-related measures of the SNS. Future studies should examine PEP in addition to electrodermal measures of galvanic skin response to systematically examine individual differences in reward-related and threat-related behavior and physiological reactivity within the same experimental context.

Future research should extend the present methodology to a wider array of experimental tasks, and examine more dynamic measures of HF-HRV and PEP. ERPs in the present study are averaged from a relatively large number of stimulus events that are largely overlapping in time, which prohibits looking at single trial dynamics that have been shown to be sensitive to associations between ERPs and cardiac physiology (Mueller, Stemmler, & Wacker, 2010). Tasks measuring ERPs during response inhibition such as the Go/No-Go and Stop Signal tasks, provide an ideal opportunity to examine whether HF-HRV and PEP interact with brain activity on the scale of milliseconds (Hajcak et al., 2003), as opposed to associating broadly at the task level. Future research will also examine growth models of HF-HRV and PEP across the four blocks of the selective attention task and the extent to which growth dynamics are associated with ERP attention effects.

In summary, these results implicate both PNS and SNS activity in individual differences in neural mechanisms of auditory selective attention. Adults with higher resting levels of HF-HRV and shorter resting levels of PEP showed larger effects of selective attention at the N1 component. While the generalizability of these findings should be interpreted with caution given the socioeconomic nature of the sample, results suggest that more efficient selective attention function is associated with heightened PNS activity and reward-related SNS activity. Given that we observe individual differences in autonomic physiology to be associated with brain activity on a task designed to control for individual differences in arousal, this raises the possibility that a variety of group-level effects commonly reported in cognitive studies would also show interactions with individual participants’ physiological state. While the present findings advocate for the inclusion of SNS measures alongside PNS measures of HF-HRV, future studies should specifically examine where electrodermal measures of SNS activity derived from skin conductance and cardiac measures of SNS activity indexed by PEP make unique contributions to neurocognitive processes. In summary, these results have the potential to inform evidence-based interventions targeting the improvement of attention within the family context, and suggest that efforts to improve attentional skills should consider how practices may alter peripheral levels of autonomic activity in addition to neural mechanisms of attention.

## Supplemental Results

Due to the small number of males (n=6) in the overall sample, results reported in the main manuscript are replicated here with females only (N=87). Any differences in findings between the two analyses are summarized in a final section, at the end of the results below.

### Characterizing physiological reactivity to the selective attention task

Paired-sample *t*-tests of baseline and task values for HF-HRV, PEP, heart rate, and respiration rate showed significant task reactivity for all measures. HF-HRV power declined from baseline (*M* = 6.12, *SD* = 1.12) to task (*M* = 6.00, *SD* = 1.08), *t*(86) = 2.44, *p* = .017. PEP values shortened from baseline (*M* = 113.57, *SD* = 10.08) to task (*M* = 112.40, *SD* = 10.28), *t*(86) = 2.45, *p* = .016. These changes in HF-HRV and PEP were associated with concurrent slowing of heart rate from baseline (*M* = 73.61, *SD* = 10.39) to task (*M* = 72.27, *SD* = 9.80), *t*(86) = 3.22, *p* = .002, and acceleration of respiration rate from baseline (*M* = 15.82, *SD* = 1.84) to task (*M* = 17.33, *SD* = 2.35), *t*(86) = -7.54, *p* < .001.

### Characterizing effects of selective attention on ERPs

For P1 mean amplitudes, results revealed an interaction of attention × laterality, *F*(2, 172) = 3.95, *p* = .023, such that significant effects of attention were seen at right-lateralized electrode clusters (*p* = .033) but not at left-lateralized or midline clusters (*ps* > .57). Follow-up comparisons showed that significant effects of attention on P1 amplitudes at the group level were seen at the right medial (*p* = .019) and right posterior (*p* = .047) electrode clusters.

For N1 mean amplitudes, results revealed a main effect of attention, *F*(1, 86) = 8.06, *p* = .006, as well as an interaction of attention × laterality, *F*(2, 172) = 12.61, *p* < .001, such that significant effects of attention were seen at left-lateralized (*p* = .002) and midline electrode clusters (*p* < .001) but not at the right-lateralized clusters (*p* = .588). Follow-up comparisons demonstrated significant attention effects on N1 amplitudes broadly across the scalp [left anterior, *p* = .015; central anterior, *p* = .012; left central, *p* = .024; central midline, *p* = .001; left posterior, *p* = .001; midline posterior, *p* < .001].

### Associations between ANS and ERP effects of selective attention

Correlations amongst all variables of interest for subsequent analyses are shown in Supplemental Table 1. Below, these results and follow-up analyses are presented separately for each ERP component. Notably, there were no significant associations between socioeconoimc risk factors and any of the autonomic or ERP measures. Therefore, socioeconomic risk was not included in the following analyses.

### P1 component (100-150 ms)

No significant associations were observed between effects of attention on P1 amplitudes and measures of HF-HRV [baseline HF-HRV: *r*(84) = .08, *p* = .478; task HF-HRV: *r*(84) = .02, *p* = .885; HF-HRV reactivity: *r*(84) = -.14, *p* = .199] or measures of PEP [baseline PEP: *r*(84) = .07, *p* = .525; task PEP: *r*(84) = .03, *p* = .818; PEP reactivity: *r*(84) = -.10, *p* = .364].

### N1 component (175-225 ms)

A significant association was observed between effects of selective attention on N1 amplitudes and baseline PEP, *r*(84) = .23, *p* = .035, with a similar effect observed between N1 amplitudes and task values of PEP, *r*(84) = .27, *p* = .014. There was a marginal trend of N1 amplitudes correlating with baseline HF-HRV, *r*(84) = -.21, *p* = .058. Shorter baseline PEP and higher baseline HF-HRV were associated with a larger effect of selective attention on N1 amplitudes. Notably, the directionality of these effects remained when controlling for respiration rate [baseline PEP, *r*(83) = .22, *p* = .044; baseline HF-HRV, *r*(83) = -.20, *p* = .072]^3^. Followup analyses of attended and unattended ERPs separately suggested that the relationship between baseline physiology and the N1 attention effect was driven by associations specific to attended ERPs. Shorter baseline PEP was associated with larger negative amplitudes to attended ERPs, *r*(83) = .23, *p* = .034, and although higher baseline HF-HRV was associated with larger negative amplitudes to attended ERPs, this relationship was non-significant, *r*(83) = -.18, *p* = .106. There was no evidence of associations between baseline physiology and unattended ERPs (*ps* > .85).

Linear regression models were then run to clarify the join contributions of baseline HF-HRV and baseline PEP with effects of selective attention on N1 amplitudes, while controlling for reactivity values and age. As shown in Supplemental Table 2, the effect of attention on N1 amplitudes was significantly predicted by baseline HF-HRV and baseline PEP [*R*^2^ = .14, *F*(3, 83) = 3.62, *p* = .016], with unique variance accounted for by baseline HF-HRV [*β* = -.23, *p* = .045] and baseline PEP [*β* = .23, *p* = .027]. Adding reactivity values of HF-HRV and PEP did not contribute additional explained variance to the model [*R*^2^ change = .02, *F* change(2, 81) = .96, *p* = .387]. To test for interactions between baseline HF-HRV and PEP, an additional model was performed including an interaction term of baseline HF-HRV x baseline PEP, but adding the interaction term did not contribute additional explained variance [*R*^2^ change = .03, *F* change(1, 80) = 2.50, *p* = .118].

### Summary of analyses after excluding male participants

Overall, there was a high degree of overlap between results for the full sample (N=93) and results for the sample when only females were considered (N=87). The same pattern of physiological reactivity was observed across HF-HRV, PEP, heart rate, and respiration rate, with very similar raw values for each measure between the two samples. Analyses of the attention effect at the P1 and N1 components were nearly identical, with the lone exception being an additional electrode cluster showing a significant N1 attention effect with the female-only sample (left central, *p* = .024), which was only marginally associated in the full sample (left central, *p* = .058).

Associations between the ERP attention effect, HF-HRV, and PEP were also very similar to analyses with the full sample. Both analyses showed no relationship between HF-HRV or PEP with the P1 attention effect, while baseline HF-HRV and PEP showed associations with the N1 attention effect. Although the association between baseline HF-HRV and the N1 attention effect was relatively attenuated in the female-only sample, *r*(84) = -.21, *p* = .058, relative to the full sample, *r*(90) = -.22, *p* = .037, when entering both baseline HF-HRV and baseline PEP into the model together with age, baseline HF-HRV remained a significant predictor of the N1 attention effect along with baseline PEP.

**Supplemental Table 1.** Partial Correlations Controlling for Age for Female Participants (N=87).

|  | 1 | 2 | 3 | 4 | 5 | 6 | 7 | 8 | 9 | 10 | 11 | 12 |
| --- | --- | --- | --- | --- | --- | --- | --- | --- | --- | --- | --- | --- |
| 1. Socioeconomic risk | - |  |  |  |  |  |  |  |  |  |  |  |
| 2. HF-HRV baseline | .06 | - |  |  |  |  |  |  |  |  |  |  |
| 3. HF-HRV task | -.03 | .88*** | - |  |  |  |  |  |  |  |  |  |
| 4. HF-HRV reactivity | -.17 | -.34** | .15 | - |  |  |  |  |  |  |  |  |
| 5. PEP baseline | -.03 | .06 | .11 | .09 | - |  |  |  |  |  |  |  |
| 6. PEP task | -.09 | .04 | .07 | .07 | .90*** | - |  |  |  |  |  |  |
| 7. PEP reactivity | -.13 | -.05 | -.08 | -.06 | -.30** | .16 | - |  |  |  |  |  |
| 8. Attended P1 amp. | .06 | -.05 | -.11 | -.10 | .17 | .07 | -.22* | - |  |  |  |  |
| 9. Unattended P1 amp. | .17 | -.15 | -.15 | .03 | .08 | .04 | -.09 | .32** | - |  |  |  |
| 10. Att. – Unatt. P1 amp. | -.09 | .08 | .02 | -.12 | .08 | .03 | -.12 | .62*** | -.54*** | - |  |  |
| 11. Attended N1 amp. | .04 | -.19t | -.16 | .08 | .25* | .29** | .07 | .03 | <.01 | .03 | - |  |
| 12. Unattended N1 amp. | .10 | .03 | .02 | -.02 | .01 | -.02 | -.02 | -.16 | .20t | -.30** | .46*** | - |
| 13. Att.- Unatt. N1 amp. | -.06 | -.22* | -.18 | .09 | .24* | .31** | .08 | .17 | -.18 | .30** | .57*** | -.46*** |

\*\*\*, $p < .001$ ; \*\*, $p < .01$ ; \*, $p < .05$ ; t, $p < .07$

**Supplemental Table 2.**
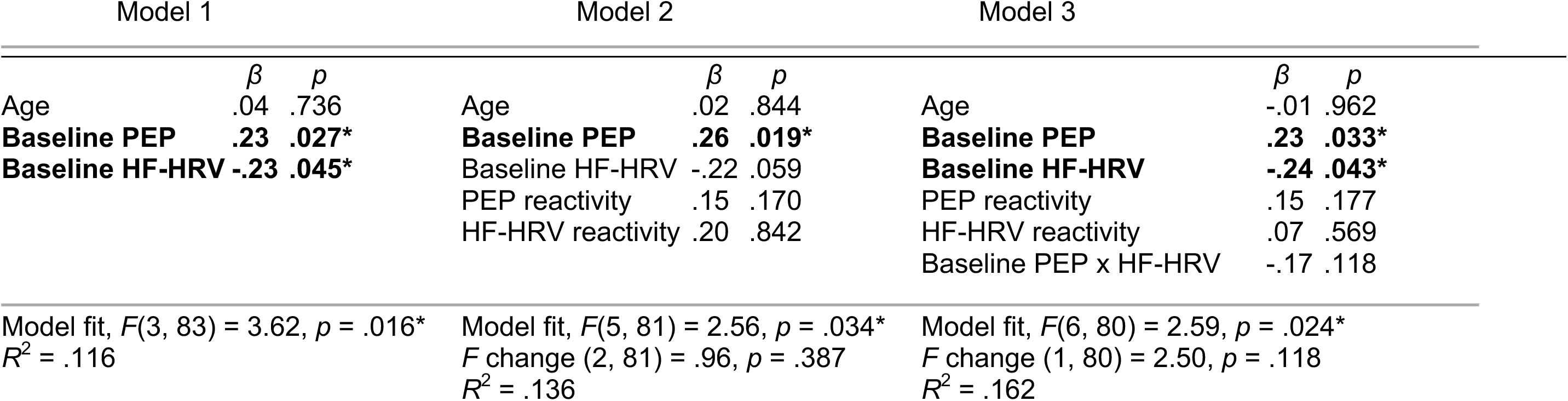
Regression Models Predicting the Effect of Selective Attention on N1 Amplitudes from Baseline HF-HRV and PEP for Female Participants (N=87).

| Model 1 |  |  | Model 2 |  |  | Model 3 |  |  |
| --- | --- | --- | --- | --- | --- | --- | --- | --- |
| | $\beta$ | $p$ | | $\beta$ | $p$ | | $\beta$ | $p$ |
| Age | .04 | .736 | Age | .02 | .844 | Age | -.01 | .962 |
| **Baseline PEP** | **.23** | **.027\*** | **Baseline PEP** | **.26** | **.019\*** | **Baseline PEP** | **.23** | **.033\*** |
| **Baseline HF-HRV** | **-.23** | **.045\*** | Baseline HF-HRV | -.22 | .059 | **Baseline HF-HRV** | **-.24** | **.043\*** |
|  |  |  | PEP reactivity | .15 | .170 | PEP reactivity | .15 | .177 |
|  |  |  | HF-HRV reactivity | .20 | .842 | HF-HRV reactivity | .07 | .569 |
|  |  |  |  |  |  | Baseline PEP x HF-HRV | -.17 | .118 |
| Model fit, $F(3, 83) = 3.62$, $p = .016^*$<br>$R^2 = .116$ | | | Model fit, $F(5, 81) = 2.56$, $p = .034^*$<br>$F$ change (2, 81) = .96, $p = .387$<br>$R^2 = .136$ | | | Model fit, $F(6, 80) = 2.59$, $p = .024^*$<br>$F$ change (1, 80) = 2.50, $p = .118$<br>$R^2 = .162$ | | |

## Footnotes

1 This analysis plan was preregistered on 05/17/16 (https://osf.io/xqncj/).

2 Effects of selective attention on N1 amplitudes were not significantly associated with baseline heart rate (*p* = .715), heart rate reactivity (*p* = .592), baseline respiration rate (*p* = .826), or respiration rate reactivity (*p* = .440).

3 Effects of selective attention on N1 amplitudes were not significantly associated with baseline heart rate (*p* = .437), heart rate reactivity (*p* = .553), baseline respiration rate (*p* = .814), or respiration rate reactivity (*p* = .397).

## References

Alderman, B. L., & Olson, R. L. (2014). The relation of aerobic fitness to cognitive control and heart rate variability: A neurovisceral integration study. Biological psychology, 99, 26-33. doi:10.1016/j.biopsycho.2014.02.007

Allen, M. T., & Crowell, M. D. (1989). Patterns of autonomic response during laboratory stressors. Psychophysiology, 26(5), 603-614. doi:10.1111/j.1469-8986.1989.tb00718.x

Allen, B., Jennings, J. R., Gianaros, P. J., Thayer, J. F., & Manuck, S. B. (2015). Resting high-frequency heart rate variability is related to resting brain perfusion. Psychophysiology, 52, 277-287. doi:10.1111/psyp.12321

Backs, R. W., & Seljos, K. A. (1994). Metabolic and cardiorespiratory measures of mental effort: the effects of level of difficulty in a working memory task. International journal of psychophysiology, 16(1), 57-68. doi:10.1016/0167-8760(94)90042-6

Beauchaine, T. (2001). Vagal tone, development, and Gray’s motivational theory: Toward an integrated model of autonomic nervous system functioning in psychopathology. Development and psychopathology, 13(02), 183-214. doi:10.1017/S0954579401002012

Beauchaine, T. P., & Thayer, J. F. (2015). Heart rate variability as a transdiagnostic biomarker of psychopathology. International journal of psychophysiology, 98(2), 338-350. doi:http://dx.doi.org/10.1016/j.ijpsycho.2015.08.004

Beissner, F., Meissner, K., Bär, K. J., & Napadow, V. (2013). The autonomic brain: an activation likelihood estimation meta-analysis for central processing of autonomic function. Journal of neuroscience, 33(25), 10503-10511. doi:10.1523/JNEUROSCI.1103-13.2013

Berntson, G. G., Cacioppo, J. T., & Fieldstone, A. (1996). Illusions, arithmetic, and the bidirectional modulation of vagal control of the heart. Biological psychology, 44(1), 1-17. doi:10.1016/S0301-0511(96)05197-6

Berntson, G. G., Cacioppo, J. T., & Grossman, P. (2007). Whither vagal tone. Biological psychology, 74, 295-300. doi:10.1016/j.biopsycho.2006.08.006

Berntson, G. G., Cacioppo, J. T., & Quigley, K. S. (1993a). Respiratory sinus arrhythmia: autonomic origins, physiological mechanisms, and psychophysiological implications. Psychophysiology, 30(2), 183-196. doi:10.1111/j.1469-8986.1993.tb01731.x

Berntson, G. G., Cacioppo, J. T., & Quigley, K. S. (1993b). Cardiac psychophysiology and autonomic space in humans: Empirical perspectives and conceptual implications. Psychophysiology, 114(2), 296-322. doi:10.1037/0033-2909.114.2.296

Berntson, G. G., Cacioppo, J. T., Quigley, K. S., Fabro, V. T. (1994). Autonomic space and psychophysiological response. Psychophysiology, 31, 44-61. doi:10.1111/j.1469-8986.1994.tb01024.x

Berntson, G. G., Cacioppo, J. T., Binkley, P. F., Uchino, B. N., Quigley, K. S., & Fieldstone, A. (1994b). Autonomic cardiac control. III. Psychological stress and cardiac response in autonomic space as revealed by pharmacological blockades. Psychophysiology, 31(6), 599-608. doi:10.1111/j.1469-8986.1994.tb02352.x

Berntson, G. G., Lozano, D. L., Chen, Y. J., & Cacioppo, J. T. (2004). Where to Q in PEP. Psychophysiology, 41(2), 333-337. doi:10.1111/j.1469-8986.2004.00156.x

Brenner, S. L., & Beauchaine, T. P. (2011). Pre-ejection period reactivity and psychiatric comorbidity prospectively predict substance use initiation among middle-schoolers: A pilot study. Psychophysiology, 48(11), 1588-1596. doi:10.1111/j.1469-8986.2011.01230.x

Bush, N. R., Alkon, A., Obradović, J., Stamperdahl, J., & Boyce, W. T. (2011). Differentiating challenge reactivity from psychomotor activity in studies of children’s psychophysiology: Considerations for theory and measurement. Journal of experimental child psychology, 110(1), 62-79. doi:10.1016/j.jecp.2011.03.004

Butler, E. A., Wilhelm, F. H., & Gross, J. J. (2006). Respiratory sinus arrhythmia, emotion, and emotion regulation during social interaction. Psychophysiology, 43(6), 612-622. doi:10.1111/j.1469-8986.2006.00467.x

Byrd, D. L., Reuther, E. T., McNamara, J. P. H., DeLucca, T. L., & Berg, W. K. (2014). Age differences in high frequency phasic heart rate variability and performance response to increased executive function load in three executive function tasks. Frontiers in psychology, 5, 1470. doi:10.3389/fpsyg.2014.01470

Capuana, L. J., Dywan, J., Tays, W. J., Elmers, J. L., Witherspoon, R., & Segalowitz, S. J. (2014). Factors influencing the role of cardiac autonomic regulation in the service of cognitive control. Biological psychology, 102, 88-97. doi:10.1016/j.biopsycho.2014.07.01

Coch, D., Sanders, L. D., & Neville, H. J. (2005). An event-related potential study of selective auditory attention in children and adults. Journal of cognitive neuroscience, 17(4), 605-622. doi:10.1162/0898929053467631

Cowan, N., Elliott, E. M., Saults, J. S., Morey, C. C., Mattox, S., Hismjatullina, A., & Conway, A. R. (2005). On the capacity of attention: Its estimation and its role in working memory and cognitive aptitudes. Cognitive psychology, 51(1), 42-100. doi:10.1016/j.cogpsych.2004.12.001

Dawson, M. E., Schell, A. M., & Filion, D. L. (2007). The electrodermal system. In J. T. Cacioppo, L. G. Tassinary, & G. G. Berntson (Eds.), Handbook of psychophysiology (3^rd^ ed., pp. 159-181). Cambridge (England): Cambridge University Press.

Delorme, A., & Makeig, S. (2004). EEGLAB: an open source toolbox for analysis of single-trial EEG dynamics including independent component analysis. Journal of neuroscience methods, 134(1), 9-21. doi:10.1016/j.jneumeth.2003.10.009

Derefinko, K. J., Eisenlohr-Moul, T. A., Peters, J. R., Roberts, W., Walsh, E. C., Milich, R., Lynam, D. R. (2016). Physiological response to reward and extinction predicts alcohol, marijuana, and cigarette use two years later. Drug and alcohol dependence, 163, S29-S36. doi:10.1016/j.drugalcdep.2016.01.034

Edelberg, R. (1993). Electrodermal mechanisms: A critique of the two-effector hypothesis and a proposed replacement. In Progress in electrodermal research (pp. 7-29). Springer, US. doi:10.1007/978-1-4615-2864-7_2

Evans, G. W., & Kim, P. (2013). Childhood poverty, chronic stress, self-regulation, and coping. Child development perspectives, 7(1), 43-48. doi:10.1111/cdep.12013

Fukuda, K., Vogel, E., Mayr, U., & Awh, E. (2010). Quantity, not quality: The relationship between fluid intelligence and working memory capacity. Psychonomic bulletin & review, 17(5), 673-679. doi:10.3758/17.5.673

Garon, N., Bryson, S. E., & Smith, I. M. (2008). Executive function in preschoolers: a review using an integrative framework. Psychological bulletin, 134(1), 31. doi:10.1037/0033-2909.134.1.31

Gatzke-Kopp, L. M. (2016). Diversity and representation: Key issues for psychophysiological science. Psychophysiology, 53(1), 3-13. doi:10.1111/psyp.12566

Gatzke-Kopp, L. M., Jetha, M. K., & Segalowitz, S. J. (2014). The role of resting frontal EEG asymmetry in psychopathology: Afferent or efferent filter? Developmental psychobiology, 56(1), 73-85. doi:10.1002/dev.21092

Gillie, B. L., Vasey, M. W., & Thayer, J. F. (2015). Individual differences in resting heart rate variability moderate thought suppression success. Psychophysiology, 52(9), 1149-1160. doi:10.1111/psyp.12443

Giuliano, R. J., Skowron, E. A., & Berkman, E. T. (2015). Growth models of dyadic synchrony and mother-child vagal tone in the context of parenting at-risk. Biological psychology, 105, 29-36. doi:10.1016/j.biopsycho.2014.12.009

Giuliano, R. J., Gatzke-Kopp, L. M., Roos, L. E., & Skowron, E. A. (2017). Resting sympathetic arousal moderates the association between parasympathetic reactivity and working memory performance in adults reporting high levels of life stress. Psychophysiology, 54(8), 1195-1208. doi:10.1111/psyp.12872

Giuliano, R. J., Karns, C. M., Neville, H. J., & Hillyard, S. A. (2014). Early auditory evoked potential is modulated by selective attention and related to individual differences in visual working memory capacity. Journal of cognitive neuroscience, 26(12), 2682-2690. doi:10.1162/jocn_a_00684

Hajcak, G., McDonald, N., & Simons, R. F. (2003). To err is autonomic: Error-related brain potentials, ANS activity, and post-error compensatory behavior. Psychophysiology, 40(6), 895-903. doi:10.1111/1469-8986.00107

Hajcak, G., McDonald, N., & Simons, R. F. (2004). Error-related psychophysiology and negative affect. Brain and cognition, 56(2), 189-197. doi:10.1016/j.bandc.2003.11.001

Hansen, A. L., Johnsen, B. H., & Thayer, J. F. (2003). Vagal influence on working memory and attention. International journal of psychophysiology, 48(3), 263-274. doi:10.1016/S0167-8760(03)00073-4

Hansen, A. L., Johnsen, B. H., Sollers, J. J., Stenvik, K., & Thayer, J. F. (2004). Heart rate variability and its relation to prefrontal cognitive function: the effects of training and detraining. European journal of applied physiology, 93(3), 263-272. doi:10.1007/s00421-004-1208-0

Hillyard, S. A., Hink, R. F., Schwent, V. L., & Picton, T. W. (1973). Electrical signs of selective attention in the human brain. Science, 182(4108), 177-180. doi:10.1126/science.182.4108.177

Hinnant, J. B., Erath, S. A., Tu, K. M., & El-Sheikh, M. (2016). Permissive parenting, deviant peer affiliations, and delinquent behavior in adolescence: the moderating role of sympathetic nervous system reactivity. Journal of abnormal child psychology, 44(6), 1071-1081. doi:10.1007/s10802-015-0114-8

Holzman, J. B., & Bridgett, D. J. (2017). Heart rate variability indices as bio-markers of top-down self-regulatory mechanisms: A meta-analytic review. Neuroscience & biobehavioral reviews, 74, 233-255. doi:10.1016/j.neubiorev.2016.12.032

Hugdahl, K. (1996). Cognitive influences on human autonomic nervous system function. Current opinion in neurobiology, 6(2), 252-258. doi:10.1016/S0959-4388(96)80080-8

Isbell, E., Wray, A. H., & Neville, H. J. (2015). Individual differences in neural mechanisms of selective auditory attention in preschoolers from lower socioeconomic status backgrounds: an event-related potentials study. Developmental science, 19(6), 865-880. doi:10.1111/desc.12334

Johnsen, B. H., Thayer, J. F., Laberg, J. C., Wormnes, B., Raadal, M., Skaret, E., Kvale, G., & Berg, E. (2003). Attentional and physiological characteristics of patients with dental anxiety. Journal of anxiety disorders,17(1), 75-87. doi:10.1016/S0887-6185(02)00178-0

Karns, C. M., Isbell, E., Giuliano, R. J., & Neville, H. J. (2015). Auditory attention in childhood and adolescence: An event-related potential study of spatial selective attention to one of two simultaneous stories. Developmental cognitive neuroscience, 13, 53-67. doi:10.1016/j.dcn.2015.03.001

Kimhy, D., Crowley, O. V., McKinley, P. S., Burg, M. M., Lachman, M. E., Tun, P. A., Ryff, C. D., Seeman, T. E., & Sloan, R. P. (2013). The association of cardiac vagal control and executive functioning-findings from the MIDUS study. Journal of psychiatric research, 47(5), 628-635. doi:10.1016/j.jpsychires.2013.01.018

Knight, R. T., Hillyard, S. A., Woods, D. L., & Neville, H. J. (1981). The effects of frontal cortex lesions on event-related potentials during auditory selective attention. Electroencephalography and clinical neurophysiology, 52(6), 571-582.

Kubota, Y., Sato, W., Toichi, M., Murai, T., Okada, T., Hayashi, A., & Sengoku, A. (2001). Frontal midline theta rhythm is correlated with cardiac autonomic activities during the performance of an attention demanding meditation procedure. Cognitive brain research, 11(2), 281-287. doi:10.1016/S0926-6410(00)00086-0

Lenneman, J. K., & Backs, R. W. (2009). Cardiac autonomic control during simulated driving with a concurrent verbal working memory task. Human factors, 51(3), 404-418. doi:10.1177/0018720809337716

Levy, M. N. (1990). Autonomic interactions in cardiac control. Annals of the New York Academy of Sciences, 601(1), 209-221. doi:10.1111/j.1749-6632.1990.tb37302.x

Lopez-Calderon, J., & Luck, S. J. (2014). ERPLAB: an open-source toolbox for the analysis of event-related potentials. Frontiers in human neuroscience, 8, 213. doi:10.3389/fnhum.2014.00213

Lozano, D. L., Norman, G., Knox, D., Wood, B. L., Miller, B. D., Emery, C. F., & Berntson, G. G. (2007). Where to B in dZ/dt. Psychophysiology, 44, 113-119. doi:10.1111/j.1469-8986.2006.00468.x

Mathewson, K. J., Jetha, M. K., Drmic, I. E., Bryson, S. E., Goldberg, J. O., Hall, G. B., Santesso, D. L., Segalowitz, S. J., & Schmidt, L. A. (2010). Autonomic predictors of Stroop performance in young and middle-aged adults. International journal of psychophysiology, 76(3), 123-129. doi:10.1016/j.ijpsycho.2010.02.007

Melis, C., & van Boxtel, A. (2001). Differences in autonomic physiological responses between good and poor inductive reasoners. Biological psychology, 58, 121-146. doi:10.1016/S0301-0511(01)00112-0

Melis, C., & van Boxtel, A. (2007). Autonomic physiological response patterns related to intelligence. Intelligence, 35, 471-487. doi:10.16/j.intell.2006.09.007

Mueller, E. M., Stemmler, G., & Wacker, J. (2010). Single-trial electroencephalogram predicts cardiac acceleration: a time-lagged P-correlation approach for studying neurovisceral connectivity. Neuroscience, 166(2), 491-500. doi:10.1016/j.neuroscience.2009

Neville, H. J., Stevens, C., Pakulak, E., Bell, T. A., Fanning, J., Klein, S., & Isbell, E. (2013). Family-based training program improves brain function, cognition, and behavior in lower socioeconomic status preschoolers. Proceedings of the national academy of sciences, 110(29), 12138-12143. doi:10.1073/pnas.1304437110

Öhman, A. (1979). The orienting response, attention, and learning: An information-processing perspective. The orienting reflex in humans, 443-471.

Park, G., & Thayer, J. F. (2014). From the heart to the mind: cardiac vagal tone modulates top-down and bottom-up visual perception and attention to emotional stimuli. Frontiers in psychology, 5, 278. doi:10.3389/fpsyg.2014.00278

Park, G., Vasey, M. W., Van Bavel, J. J., & Thayer, J. F. (2013). Cardiac vagal tone is correlated with selective attention to neutral distractors under load. Psychophysiology, 50(4), 398-406. doi:10.1111/psyp.12029

Park, G., Vasey, M. W., Van Bavel, J. J., & Thayer, J. F. (2014). When tonic cardiac vagal tone predicts changes in phasic vagal tone: The role of fear and perceptual load. Psychophysiology, 51(5), 419-426. doi:10.1111/psyp.12186

Porges, S. W. (2007). The polyvagal perspective. Biological psychology, 74(2), 116-143. doi:10.1016/j.biopsycho.2006.06.009

Sanders, L. D., Stevens, C., Coch, D., & Neville, H. J. (2006). Selective auditory attention in 3-to 5-year-old children: An event-related potential study. Neuropsychologia, 44(11), 2126-2138. doi:10.1016/j.neuropsychologia.2005.10.007

Saus, E. R., Johnsen, B. H., Eid, J., Risem, P. K., Anderson, R., & Thayer, J. F. (2006). The effect of brief situational awareness training in a police shooting simulator: An experimental study. Military psychology, 18(Supplemental), S3-S21. doi:10.1207/s15327876mp1803s_2

Segerstrom, S. C., & Nes, L. S. (2007). Heart rate variability reflects self-regulatory strength, effort, and fatigue. Psychological science, 18(3), 275-281. doi:10.1111/j.1467-9280.2007.01888.x

Shonkoff, J. P., & Fisher, P. A. (2013). Rethinking evidene-based practice and two-generation programs to create the future of early childhood policy. Development and psychopathology, 25, 1636-1653. doi:10.1017/S0954579413000813

Skowron, E. A., Cipriano-Essel, E., Gatzke-Kopp, L. M., Teti, D. M., & Ammerman, R. T. (2014). Early adversity, RSA, and inhibitory control: Evidence of children’s neurobiological sensitivity to social context. Developmental psychobiology, 56(5), 964-978. doi:10.1002/dev.21175

Smith, R., Thayer, J. F., Khalsa, S. S., & Lane, R. D. (2017). The hierarchical basis of neurovisceral integration. Neuroscience & biobehavioral reviews. doi:10.1016/j.neubiorev.2017.02.003

Spangler, D. P., Bell, M. A., & Deater-Deckard, K. (2015). Emotion suppression moderates the quadratic association between RSA and executive function. Psychophysiology, 52(9), 1175-1185. doi:10.1111/psyp.12451

Stevens, C., Lauinger, B., & Neville, H. (2009). Differences in the neural mechanisms of selective attention in children from different socioeconomic backgrounds: An event-related brain potential study. Developmental science, 12(4), 634-646. doi:10.1111/j.1467-7687.2009.00807.x

Thayer, J. F., Ahs, F., Fredrikson, M., Sollers III, J. J., & Wager, T. D. (2012). A meta-analysis of heart rate variability and neuroimaging studies: Implications for heart rate variability as a marker of stress and health. Neuroscience and biobehavioral reviews, 36, 747-756. doi:10.1016/j.neubiorev.2011.11.009

Thayer, J. F., & Lane, R. D. (2000). A model of neurovisceral integration in emotion regulation and dysregulation. Journal of affective disorders, 61(3), 201-216. doi:10.1016/S0165-0327(00)00338-4

Thayer, J. F., & Lane, R. D. (2009). Claude Bernard and the heart–brain connection: Further elaboration of a model of neurovisceral integration. Neuroscience & Biobehavioral Reviews, 33(2), 81-88. doi:10.1016/j.neubiorev.2008.08.004

Tranel, D., & Damasio, H. (1994). Neuroanatomical correlates of electrodermal skin conductance responses. Psychophysiology, 31(5), 427-438. doi:10.1111/j.1469-8986.1994.tb01046.x

Triggiani, A. I., Valenzano, A., Del Percio, C., Marzano, N., Soricelli, A., Petito, A., Bellomo, A., Basar, E., Mundi, C., Cibelli, G., & Babiloni, C. (2016). Resting state Rolandic mu rhythms are related to activity of sympathetic component of autonomic nervous system in healthy humans. International journal of psychophysiology, 103, 79-87. doi:10.1016/j.ijpsycho.2015.02.009

Tsukahara, J. S., Harrison, T. L., & Engle, R. W. (2016). The relationship between pupil size and intelligence. Cognitive psychology, 91, 109-123. doi:10.1016/j.cogpsych.2016.10.001

Williams, D. P., Thayer, J. F., & Koenig, J. (2016). Resting cardiac vagal tone predicts intraindividual variability during an attention task in a sample of young and healthy adults. Psychophysiology, 53(12), 1843-1851. doi:10.1111/psyp.12739

Zahn, D., Adams, J., Krohn, J., Wenzel, M., Mann, C. G., Gomille, L. K., Jacobi-Scherbening, V., & Kubiak, T. (2016). Heart rate variability and self-control – A meta-analysis. Biological psychology, 115, 9–26. doi:10.1016/j.biopsycho.2015.12.007

